# Learning Context-aware Structural Representations to Predict Antigen and Antibody Binding Interfaces

**DOI:** 10.1101/658054

**Authors:** Srivamshi Pittala, Chris Bailey-Kellogg

## Abstract

Understanding how antibodies specifically interact with their antigens can enable better drug and vaccine design, as well as provide insights into natural immunity. Experimental structural characterization can detail the “ground truth” of antibody-antigen interactions, but computational methods are required to efficiently scale to large-scale studies. In order to increase prediction accuracy as well as to provide a means to gain new biological insights into these interactions, we have developed a unified deep learning-based framework to predict binding interfaces on both antibodies and antigens. The framework leverages three key aspects of antibody-antigen interactions in order to learn predictive structural representations: (1) since interfaces are formed from multiple residues in spatial proximity, we employ graph convolutions to aggregate properties across local regions in a protein; (2) since interactions are specific between antibody-antigen pairs, we employ an attention layer to explicitly encode the context of the partner; (3) since more data is available for general protein-protein interactions, we employ transfer learning to leverage this data as a prior for the specific case of antibody-antigen interactions. We show that this single framework achieves state-of-the-art performance at predicting binding interfaces on both antibodies and antigens, and that each of its three aspects drives additional improvement in the performance. We further show that the attention layer not only improves performance, but also provides a biologically interpretable perspective into the mode of interaction.

## 1 Introduction

As one of its mechanisms to combat disease, the immune system develops B cells that secrete antibodies to specifically recognize and either neutralize or help drive functional responses against a pathogen. An antibody recognizes a particular region, called its epitope, on a particular part of the pathogen, called its antigen; the region of the antibody directly involved in the recognition is called its paratope. The interface between an epitope and paratope is crucial to the affinity and specificity of an antibody-antigen interaction, and thus the antibody’s function. Characterizing antibody-antigen interactions at the epitope-paratope resolution can thus reveal mechanisms of immune recognition, and, over a set of antibodies, can even provide insights into the development of the immune response. Such characterization can also benefit the development of therapeutics and vaccines. For example, therapeutic antibodies are being used to treat many different diseases [7, 20], and early development processes typically yield large arrays of candidate antibodies from which to select. Understanding their different recognition mechanisms can aid the selection and subsequent development. Similarly, subunit vaccines are being developed to train the immune system against a pathogen by mimicking an important part but without causing actual infection [5, 11, 12]. Understanding the recognition processes driving a beneficial response, as well as those that are not useful, can guide the development of these vaccines so as to ensure the desired immune targeting.

Experimental structure determination methods, namely x-ray crystallography, NMR spectroscopy, and cryoEM, provide the gold standard for characterizing antibody-antigen binding modes [3, 28]. Unfortunately they remain expensive and time-consuming, and cannot feasibly keep up with the exploding amount of antibody sequence data for which it is desirable to understand antigen recognition, e.g., the millions of sequences obtained from analysis of an immune repertoire [33, 50, 56]. Alternative experimental methods like H-D exchange mass spectrometry [17] and alanine scanning [53] are faster and cheaper, and of lower resolution/confidence, but still require substantial experimental effort per target. Higher-throughput methods such as multiplexed SPR can characterize many interactions simultaneously but do not provide direct localization information [38, 6]. Computational methods thus have the most promise to scale to characterization of large numbers of possible epitope-paratope interactions, but it is necessary to ensure that predictions provide sufficient grounds to support further investigations, in terms of overall accuracy as well as the underlying reasoning for a prediction. Prediction of antibody-antigen binding interfaces can be seen as a special case of predicting protein-protein binding interfaces. However, since these interfaces have their own special characteristics [27, 15] (as do other classes of protein-protein interactions), specific methods have been developed for epitope prediction and others for paratope prediction. Many methods make predictions based on amino acid sequence alone, e.g., predicting epitopes based on neural networks [39], SVMs [14, 43], HMMs [55], and random forests [21], and paratopes using LSTMs [31, 10] and random forests [34] Though sequence-based methods can perform well on paratope prediction, most sequence-based epitope predictions are limited to the special case of a sequentially contiguous epitope [54], while in contrast most epitopes are found to be conformational (distal in sequence, but close in 3D structure) [52, 37]. Thus we and others focus on structure-based methods that leverage geometric information in making predictions. Fortunately, while the complex structure is not known, in most common scenarios the structure of the antigen by itself is available, and antibody structure prediction techniques enable confident prediction of most of the antibody’s structure [44, 45]. We thus briefly review this body of most closely related work on structure-based prediction of epitopes and paratopes.

### Docking

Many structure-based methods for epitope and paratope prediction rely on computational docking techniques, which estimate the most likely conformations of a complex based on complementarity (geometric, chemical, energetic) between the individual proteins in many possible poses [8, 40, 46]. The resulting docking models may be ranked using a scoring function incorporating many different geometric and physico-chemical parameters; defining a good scoring function is a challenging task that typically relies on domain expertise [35]. From the top ranked conformations, regions on one protein that are close to the partner protein can be identified as binding interfaces. Thus antibody-antigen docking can simultaneously predict epitopes and paratopes. While the recall of docking is generally fairly high if enough docking models are considered, the precision is then fairly low, prompting the development of methods that directly predict epitopes and paratopes based on properties of the proteins [4].

### Epitope prediction

Some approaches, e.g., PEPITO [49], ElliPro [36], EPSVR [30], and DiscoTope [25], apply machine learning methods to structural features of the antigen’s residues. These methods can be considered antibody-agnostic as they do not use information from the partner antibody, and thus just reveal parts of the antigen generally amenable to antibody binding [42]. For prediction of an epitope targeted by a particular antibody, the context of which residues are likely to be involved in the interaction can improve the prediction performance, as well as distinguish specificity differences among different antibodies. This aspect of antibody-antigen interactions was leveraged by the antibody-specific prediction method EpiPred [24] to achieve state-of-the-art performance. EpiPred first performs geometric matching of patches (e.g., based on docking models) and then scores residues on the antigen with a customized binding potential specific for antibodies and antigens.

### Paratope prediction

Many paratope prediction predictors focus on special regions on antibodies called complementarity determining regions (CDRs), as they are well-defined from sequence and constitute the majority of the paratope and the majority of the differences among antibodies driving antigen-specific recognition. In the non-parametric method Paratome, the query antibody’s structure and sequence are compared against a non-redundant dataset of antibodies, and paratopes are predicted based on resemblance to those on the closest matching antibody. [26]. Antibody i-Patch [23] uses a scoring function derived from an analysis of antibody-antigen interactions in a non-redundant training set. Recently, Daberdaku et al. [9] achieved state-of-the-art performance with a method that applies SVMs to classify patches extracted from the surface of the antibody, based on roto-translationally invariant shape descriptors and other physico-chemical properties representing the patches.

A common drawback of current structure-based methods for epitope and paratope prediction is the use of fixed representations, which can be limited by the extent of available domain knowledge. Furthermore, epitope and paratope prediction are treated as two separate tasks, leading to the use of different representations and prediction methods for antigens and antibodies. Sequence-based methods Parapred [31] and AG-Parapred [10] demonstrate the utility of learning representations for better paratope prediction. However, there are currently no methods to learn structural representations for either epitope or paratope prediction tasks. Recently, a spatial graph convolution network was proposed to learn structural representations of proteins for interface prediction in general protein-protein interactions [16]. While graph convolution networks can encode structural representations of residues with information from their spatial neighborhood, they do not encode the context of the target protein. As shown by current methods, embedding the correct context of the target protein can improve the prediction performance [24, 23]. Therefore, there is a need to develop methods for learning context-aware structural representations for epitope and paratope prediction.

In this work, we present a unified deep learning-based framework for learning context-aware structural representations of antigens and antibodies to predict their binding interfaces. Our framework consists of a novel combination of graph convolution networks, attention, and transfer learning to capture several desired aspects of antibody-antigen interactions. We show that the models trained on our framework can overcome the limitations of current computational methods and achieve state-of-the-art performance on both epitope and paratope prediction tasks. Using the attention layer, we demonstrate the ability of our framework to reveal the mode of interaction between antigens and antibodies, enabling a deeper study of the biological factors driving their interactions. Therefore, our framework improves prediction accuracy and provides interpretable results to expedite the process of large-scale antibody-antigen characterization.

## 2 Learning context-aware structural representations

We propose a novel deep learning framework (Figure 1) to learn structural representations of antigens and antibodies in order to predict their binding interfaces (i.e., antigen epitopes and antibody paratopes). Our framework comprises three components to leverage biological insights: (1) graph convolutions to capture the spatial relationships of the interfaces, (2) an attention layer to enable each protein’s interface predictions to account for the potential binding context provided by its partner, and (3) transfer learning to leverage the larger set of data available for general protein-protein interactions to provide a baseline model to be fine-tuned with antibody-antigen data.

**Figure 1:**
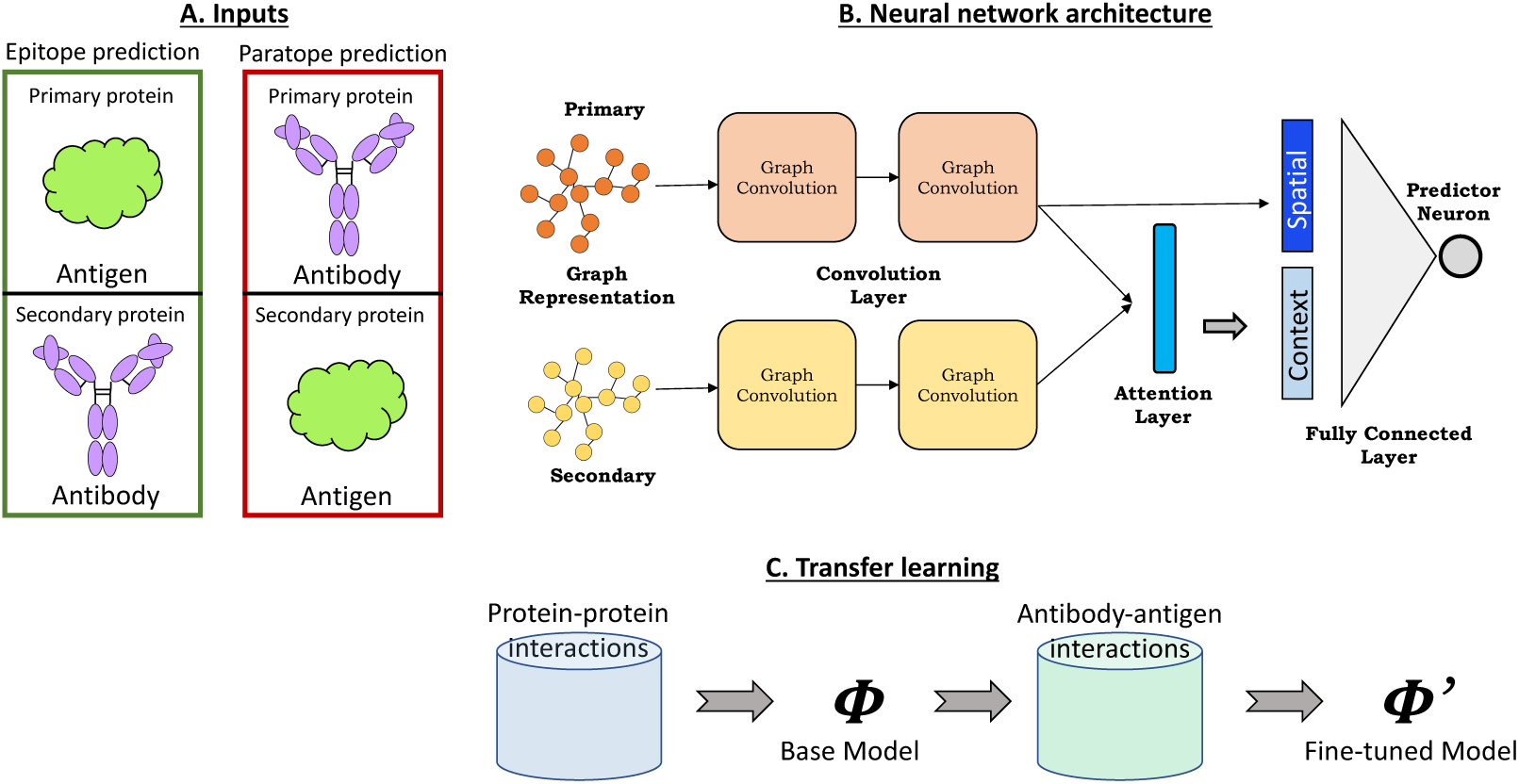
Schematic overview of the proposed framework.

We use this general framework to train two separate networks for the two prediction tasks: (1) an epitope prediction network in which the antigen is the primary protein on which we want to predict the interface (epitope) and the antibody is the secondary protein providing the context for a suitable interface; (2) a paratope prediction network with the antibody as primary for interface prediction (paratope) and the antigen secondary providing the context. We note that in both tasks, the interface labels of the secondary protein are hidden during training and prediction phases, forcing the attention layer to learn the correct context of the secondary protein in an unsupervised fashion.

### Problem statement

The objective is to assign a label, either +1 (interface) or −1 (non-interface), to each residue of the primary protein.

#### A. Input representation

Each protein structure is represented as a graph, with nodes for the amino acid residues and edges between residues with C*β*-C*β* distance less than 10Å. Associated with each node is a 62-dimension feature vector encoding important sequence and structural properties as used in [16]: (a) a one-hot encoding of the amino acid type (*d*=20); (b) a conservation profile for that position across a set of homologous proteins returned by PSI-BLAST [2] (*d*=20); (c) the absolute and relative solvent accessible surface area of the residue as computed by STRIDE [19] (*d*=2); (d) a local amino acid profile indicating the number of times each amino acid type appears within 8Å of the residue (*d*=20). The structure-based features (c and d) were calculated for each protein in isolation from its partner.

Since antibody CDRs drive their antigen-specific recognition and the rest of the antibody framework is quite similar across all antibodies, nodes in the antibody graph are limited to “CDR clouds” as follows: (1) identify the six CDRs using the IMGT annotation tool [29]; (2) for each CDR, consider two sequentially adjacent residues; (3) further extend these sets to include all residues within 6Å in the structure (the maximum of the minimum C*β*-C*β* distance between any two CDR residues in the training sets).

#### B. Neural network

The neural network consists of graph convolution, attention, and fully connected layers. Given two input graphs, primary 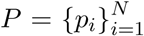 and secondary 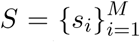, the network assigns to each node *p*_*i*_ ∈*P* a probability of belonging to the positive class (i.e., binding interface).

##### B.1 Graph convolutions

Graph convolution [16] enables order-independent aggregation of properties over a neighborhood of residues that together contribute to the formation of a binding interface. For a node *x*_*i*_ and its receptive field consisting of *K* spatial neighbors 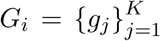 from the input graph, the convolution operation results in a vector 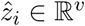, where *v* is a specified number of filters for the layer (Eqn. 1). The parameters of this operation represent the aggregation weight matrix **W**^**c**^ for the center node, the aggregation weight matrix **W**^**g**^ for the neighboring nodes, and the bias vector 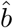. Thus, the convolution operation for a node *x*_*i*_ results in a spatial vector representation 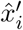 in the latent space ℝ^*v*^.

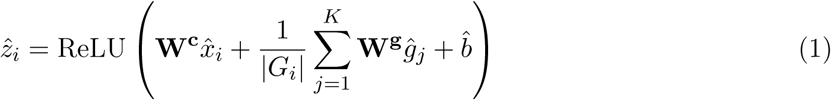

Multiple layers can be stacked to produce high-level representations for each node. Each convolution layer has two weight-shared graph convolution modules, one for the primary graph and one for the secondary graph.

##### B.2 Attention

An attention layer encodes the context of secondary graph in the residue-level representations of the primary graph, providing information for each primary residue about secondary residues that are likely to interact with it. An attention score *a*_*ij*_ is computed between all node pairs *p*_*i*_ ∈*P* and *s*_*j*_ ∈*S* after projecting them into a latent space via an attention weight matrix **W**^**a**^ (Eqn. 2). The dimensions of **W**^**a**^ are determined by the number of neurons in the final convolution layer and the desired dimension of the latent space. This dot product style of computing attention scores was used to directly estimate complementarity between hidden representations as in [32, 10]. The context vector 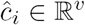 for node *p*_*i*_ is then computed by aggregating the node-level representations of *S* using normalized attention scores (Eqn. 3). The normalization was performed to calibrate the score between each pair with respect to scores across all possible pairs. The normalized score *α*_*ij*_ can therefore be interpreted as a pair-wise interaction potential between *p*_*i*_ and *s*_*j*_.

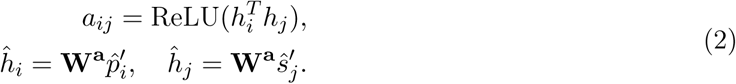

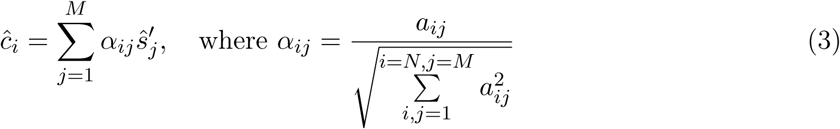

##### B.3 Node classification

A final fully-connected layer performs classification for each primary node *p*_*i*_ based on its spatial vector 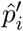 and context vector 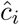 (Eqn. 4). A logit function transforms each node’s output *y*_*i*_ to indicate the probability of belonging to the positive class.

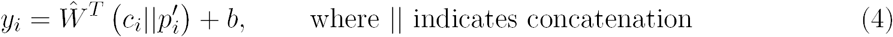

#### C. Transfer learning

A base network *ϕ* for interface prediction is learned for a relatively larger set of general protein-protein interactions. The learned weights from the base model are then used to initialize weights for training the two task-specific networks, essentially fine-tuning the general base network for epitope and paratope prediction using antibody-antigen data.

## 3 Experiments

We evaluate our approach in head-to-head benchmark comparisons against state-of-the-art epitope and paratope predictors, showing that our unified framework outperforms approaches specifically targeted to each. Furthermore, we elaborate general precision and recall trends in the full architecture, as well as versions enabling characterization of the contributions of convolution, attention, and transfer learning. Finally, we explore the ability of the attention layer to provide insights into the basis for the model’s predictions.

### 3.1 Datasets

#### Epitope prediction

The dataset from EpiPred [24] consists of 148 antibody-antigen complexes, 118 for training and 30 for testing. Since a separate validation set was not used, we constructed one from the antibody-antigen complexes in the Docking Benchmarking Dataset (DBD) v5 [51], ensuring that there were no overlapping complexes (by PDB id) among the three sets. This yielded resulted in 103 complexes for training, 38 for validation, and 30 for testing.

#### Paratope prediction

The dataset from Daberdaku et al. [9] consists of 471 antibody-antigen complexes, with 213 complexes for training, 106 for validation, and 152 for testing. Since our framework accepts only proteins, we discarded complexes with non-protein antigens (e.g., DNA), resulting in 205 complexes for training, 103 for validation, and 152 for testing.

#### Transfer learning

To facilitate unbiased transfer learning, the DBD v5 dataset [51] was processed to discard complexes that were categorized as antibody-antigen, resulting in a dataset of 191 protein-protein complexes.

Following previous studies [24, 9], residues were labeled as part of the interface if they had any non-Hydrogen atoms within 4.5Å of any non-Hydrogen atoms of residues on the other protein.

Table1 summarizes these dataset characteristics.

**Table 1:** Summary of datasets used for training, validation, and testing.

| Epitope prediction dataset |  |  | Paratope prediction dataset |  |  |
| --- | --- | --- | --- | --- | --- |
| Data split | # Complexes | % Epitopes | Data split | # Complexes | % Paratopes |
| Training | 103 | 8.9% | Training | 205 | 8.8% |
| Validation | 38 | 9.7% | Validation | 103 | 8.9% |
| Test | 30 | 7.8% | Test | 152 | 9.4% |

### 3.2 Implementation Details

The framework was implemented in TensorFlow [1]. The validation sets were used to find the optimal set of network training parameters for final evaluation. A grid search was performed over the following parameters: (a) Optimizer: Stochastic gradient descent, Momentum [48], Adagrad [13], or Adam [22]]; (b) learning rates: 0.0001, 0.001, 0.005, 0.01, 0.05, or 0.1; (c) batch size: 32, 64, pr 128; (d) dropout: 0.5 or 0.8. For each combination, networks were trained until the performance on validation set stopped improving or for a maximum of 250 epochs. For both epitope and paratope prediction, the best validation set performance was achieved when training till 120 epochs using the Momentum optimizer with Nesterov accelerated gradients [47] at a learning rate of 0.001, with batch size of 32 and 50% dropout rate. Training was carried out by minimizing the weighted cross-entropy loss function as in [16]. The same network settings were used for training on general protein-protein complexes, but the fine-tuning was carried out on antibody-antigen complexes for half the original time (i.e., 60 epochs). The graph convolution layers were set to have 32 filters and the latent space dimension for attention was also set to 32. All weight matrices were initialized as in [18] and biases set to zero. For graph convolution, the receptive field (spatial neighborhood) for each node was set to include the 15 nearest nodes in the graph.

### 3.3 Evaluation

Epitope and paratope prediction networks were trained using the validation-set optimized hyper-parameters from above. Per-protein prediction performance was measured on the test sets by comparing predicted scores against ground truth labels. While our networks output a probability for each residue, in order to enable a direct comparison to other epitope prediction methods we computed precision and recall by predicting as interface residues those with logits above 0.5. To further elaborate precision-recall trade-offs, we considered all such classification thresholds and computed the area under the precision recall curve (AUC-PR). Though some previous methods have used the AUC-ROC metric, AUC-PR is more suitable here since the emphasis is on predicting binding interfaces (positive class) and the negative class constitutes roughly 90% of the samples. To summarize the performance, AUC-PR was averaged over all proteins in the test set. To provide robust estimates of performance, the training and testing procedures were repeated five times, and the mean and standard error reported. Our evaluations included two learning schema: task-specific learning (i.e., just using antibody-antigen data) and transfer learning (i.e., fine-tuning from a model trained with general protein-protein data). For each schema, five networks were evaluated: one network with a single fully-connected layer (No convolution), one with a single graph convolution layer (Conv1-layer) and one with two (Conv2-layer), and likewise one network with the attention layer following a single graph convolution layer (Conv1-layer+Attn) and one following two convolution layers (Conv2-layer+Attn).

### 3.4 Results

#### Epitope prediction

Table 2 summarizes the epitope test set prediction performance for our different neural network implementations along with the state-of-the-art Epipred [24] and DiscoTope [25]. Our networks all perform better than Epipred and DiscoTope in terms of precision and recall at the 0.5 cutoff. Elaborating performance for a range of cut-offs via AUC-PR enables further comparison among our architecture implementations (these numbers are not available for the other methods). The network with an attention layer after two convolution layers achieves the best performance, confirming the utility of embedding the context of the target antibody into the representation of antigen’s residues in addition to information from their spatial neighbors. Furthermore, the improvements in performance of all models after transfer learning illustrates the benefits of leveraging data from general protein-protein interactions to establish a base model that can be fine-tuned with antibody-antigen data.

**Table 2:** Epitope prediction performance summary. The measures for EpiPred and DiscoTope were taken from [24]

| Method | Task-specific learning |  |  | Transfer learning |  |  |
| --- | --- | --- | --- | --- | --- | --- |
|  | AUC-PR | Precision | Recall | AUC-PR | Precision | Recall |
| No Convolution | 0.177±0.035 | 0.153 | 0.601 | 0.191±0.021 | 0.162 | 0.500 |
| Conv1-layer | 0.193±0.005 | 0.152 | 0.725 | 0.191±0.003 | 0.160 | 0.582 |
| Conv1-layer+Attn | 0.187±0.004 | 0.147 | 0.683 | 0.202±0.002 | 0.160 | 0.580 |
| Conv2-layer | 0.192±0.005 | 0.147 | 0.737 | 0.226±0.004 | 0.157 | 0.678 |
| Conv2-layer+Attn | **0.212±0.007** | 0.154 | 0.691 | **0.242±0.006** | 0.165 | 0.579 |
| Epipred [24] | NA | 0.136 | 0.436 | - | - | - |
| DiscoTope [25] | NA | 0.214 | 0.110 | - | - | - |

#### Paratope prediction

Table 3 summarizes the paratope test set prediction performance of our different neural networks and state-of-the-art structure-based methods Daberdaku et al. [9] and Antibody i-Patch [23]. To enable a direct comparison to previous studies, we predict for the entire structure of the antibody Fv region instead of just the CDR clouds as described in our methods. Our networks perform better than the other methods on both AUC-PR and AUC-ROC, establishing the superior performance of learned features over pre-defined features as used by Daberdaku et al. The network with a single layer of convolution and attention achieves the best performance, but the attention layer provides only a small performance improvement over convolution. We hypothesize that since paratopes are mostly localized to regions around the CDRs, the context of the antigen may not provide much more information regarding exact paratope location than the structural properties already captured by convolution. Nonetheless, as we show in the next section, the attention layer offers the benefit of making the network interpretable, which can be a difficult task for convolution layers alone.

**Table 3:** Paratope prediction performance summary. The performance for Daberdaku et al.[9] and Antibody-iPatch[23] were taken from [9]

| Method | Task-specific learning |  | Transfer learning |  |
| --- | --- | --- | --- | --- |
|  | AUC-PR | AUC-ROC | AUC-PR | AUC-ROC |
| No Convolution | 0.652±0.012 | 0.938±0.004 | 0.658±0.003 | 0.937±0.001 |
| Conv1-layer | **0.700±0.002** | 0.956±0.000 | 0.696±0.001 | 0.957±0.000 |
| Conv1-layer+Attn | **0.700±0.001** | **0.957±0.000** | **0.703±0.000** | **0.958±0.001** |
| Conv2-layer | 0.691±0.004 | 0.957±0.000 | 0.689±0.002 | 0.957±0.000 |
| Conv2-layer+Attn | 0.692±0.004 | 0.957±0.001 | 0.697±0.001 | 0.958±0.000 |
| Daberdaku et al.[9] | 0.658 | 0.950 | - | - |
| Antibody i-Patch[23] | 0.376 | 0.840 | - | - |

**Table 4:** Performance summary from “cross-task evaluation”, predicting paratopes using networks trained only for predicting epitopes.

| Method | Cross-task evaluation |  | Task-specific learning |  |
| --- | --- | --- | --- | --- |
|  | AUC-PR | AUC-ROC | AUC-PR | AUC-ROC |
| Conv1-layer | 0.612±0.028 | 0.844±0.026 | 0.770±0.000 | 0.970±0.007 |
| Conv1-layer+Attn | 0.624±0.026 | 0.875±0.035 | 0.768±0.003 | 0.971±0.008 |
| Conv2-layer | 0.588±0.058 | 0.910±0.051 | 0.766±0.005 | 0.971±0.001 |
| Conv2-layer+Attn | **0.628±0.025** | **0.938±0.029** | 0.765±0.007 | 0.971±0.012 |

#### Assessing the contributions of attention

The attention layer provides the opportunity to study the mode of interaction by revealing the learned context of the target protein without requiring additional inference techniques. The attention score between every pair of residues can be visualized as a matrix, as Figure 2A illustrates for the complex on which our epitope prediction network performed best. In this heatmap, epitopes have a substantially distinct attention profile compared to other residues on the antigen, which results in improved epitope prediction (AUC-PR: 0.6) compared to convolution alone (AUC-PR: 0.45). These attention scores can further be projected onto the structures (Figure 2B) by taking for each residues the maximum of its scores with partner residues. This projection shows that attention is high between residues in and around the actual interface region, suggesting that the attention layer encodes the correct context (i.e., paratopes) of the antibody for epitopes. The same pattern of high attention scores near the interface regions was also observed for other antibody-antigen complexes (Figure. 2C-F illustrates the next 4 top performers).

**Figure 2:**
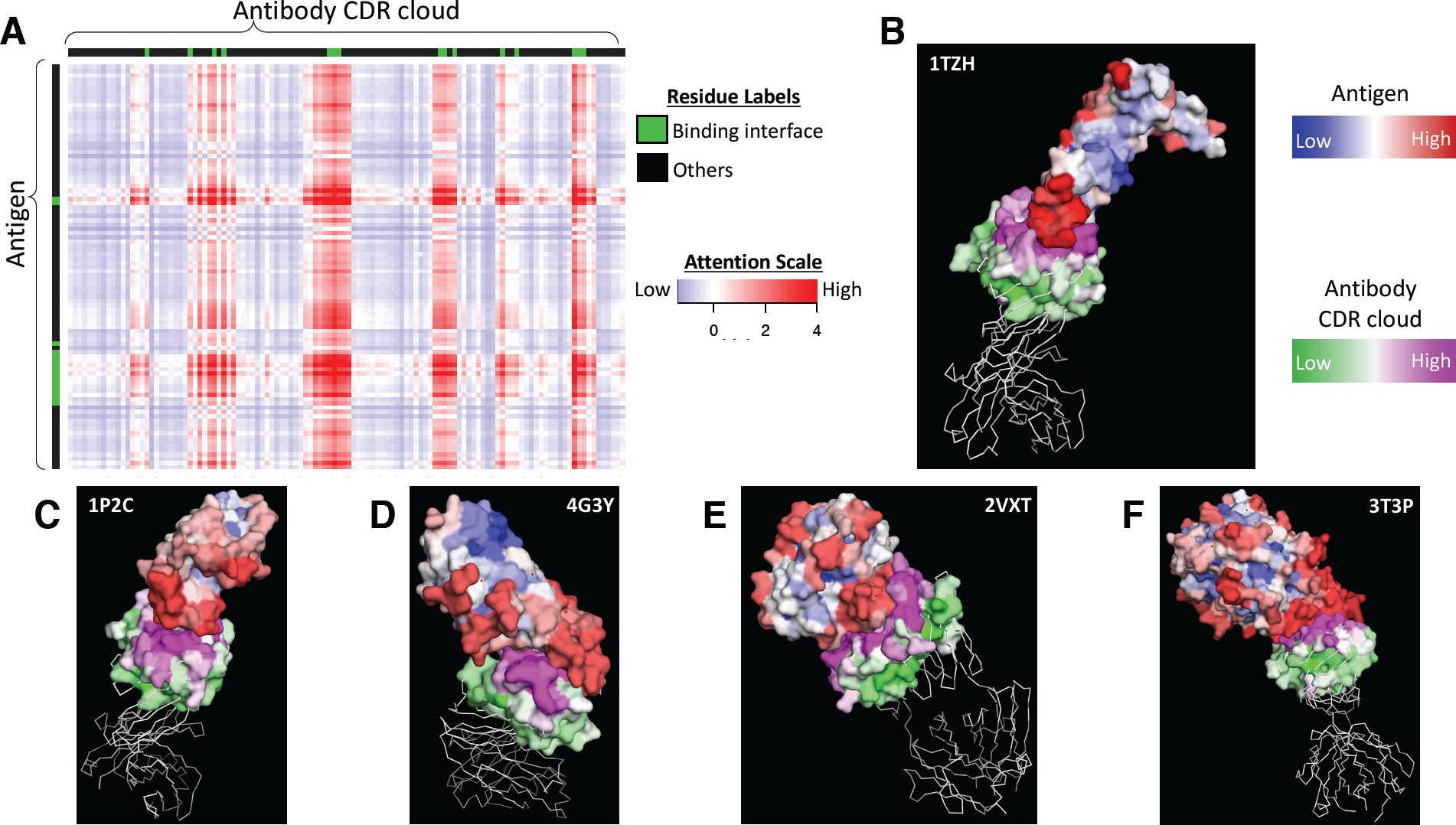
Attention visualization (A) Heatmap of the attention score matrix for an antibody in complex with VEGFA (vascular endothelial growth factor A) (PDB ID 1TZH). The scores were normalized to have zero mean and unit variance, and truncated to the range [−4,4]. (B-E) Projection of max-pooled attention scores onto structures of example antigen and antibody complexes in the test set. Structural visualizations made using PyMOL [41].

Intrigued by the attention layer’s ability to localize the appropriate context during epitope prediction, we hypothesized that the same ability could benefit paratope prediction. We thus performed a “cross-task evaluation”, in which a network was trained to predict *epitopes* using the *antigens* in the epitope prediction training set. This epitope prediction network was then evaluated for its performance at also predicting *paratopes* for the *antibodies* in the epitope prediction test set—the reciprocal task to that for which it was trained. For reference, paratope prediction networks were trained on the antibodies from the epitope prediction training set and applied (as normal) to the antibodies from the epitope prediction test set. As expected, the paratope prediction networks perform significantly better at predicting epitopes than do the networks trained to predict epitopes. However, the results from cross-task evaluation show that even though none of the networks were trained to predict paratopes, those with an attention layer perform better than convolution-only networks. This suggests that the attention layer is indeed able to better capture the specificity of antibody-antigen interactions, thereby also benefiting paratope prediction.

## 4 Conclusion

We have presented a unified deep learning framework for predicting binding interfaces on antibodies and antigens. Our results demonstrate that the networks learn structural representations that capture many desired aspects of antibody-antigen interactions and simultaneously achieve state-of-the-art performance on both epitope and paratope prediction tasks. We also show that the attention layer successfully encodes the context of partner proteins, improving prediction performance and providing an interpretable view of the mode of interaction. Future work includes including additional residue features while imposing sparsity constraints on the attention matrix, applying the same framework to other large protein families with specific recognition modes, and using predictions to focus docking as well as experimental evaluation.

## Acknowledgments

This research was supported in part by NIH grant 2R01GM098977, with computational resources from NSF award CNS-1205521 and discovery clusters at Dartmouth. We thank Margaret Ackerman, Karl Griswold, and members of the Bailey-Kellogg lab for helpful comments, and Casey Hua for sharing antibody art in Figure 1.

## Code availability

The source code will be made available upon publication of the manuscript.

